# Competitive state of actions during planning predicts sequence execution accuracy

**DOI:** 10.1101/2020.05.08.085068

**Authors:** Myrto Mantziara, Tsvetoslav Ivanov, George Houghton, Katja Kornysheva

**Affiliations:** School of Psychology, Bangor University, Bangor, Wales LL57 2AS, UK; Bangor Imaging Unit, Bangor University, Bangor, Wales LL57 2AS, UK

**Keywords:** motor sequence, preparation, reaction time, finger accuracy, competitive queuing

## Abstract

Humans can learn and retrieve novel skilled movement sequences from memory, yet the content and structure of sequence planning are not well understood. Previous computational and neurophysiological work suggests that actions in a sequence are planned as parallel graded activations and selected for output through competition (competitive queuing; CQ). However, the relevance of CQ during planning to sequence fluency and accuracy, as opposed to sequence timing, is unclear. To resolve this question, we assessed the competitive state of constituent actions behaviourally during sequence preparation. In three separate multi-session experiments, 55 healthy participants were trained to retrieve and produce 4-finger sequences with particular timing from long-term memory. In addition to sequence production, we evaluated reaction time (RT) and error rate increase to constituent action probes at several points during the preparation period. Our results demonstrate that longer preparation time produces a steeper CQ activation and selection gradient between adjacent sequence elements, whilst no effect was found for sequence speed or temporal structure. Further, participants with a steeper CQ gradient tended to produce correct sequences faster and with a higher temporal accuracy. In a computational model, we hypothesize that the CQ gradient during planning is driven by the width of acquired positional tuning of each sequential item, independently of timing. Our results suggest that competitive activation during sequence planning is established gradually during sequence planning and predicts sequence fluency and accuracy, rather than the speed or temporal structure of the motor sequence.

**Highlights:**

- Pre-ordering of actions during sequence planning can be assessed behaviourally
- Competitive gradient reflects sequence preparedness and skill, but not speed or timing
- Gradient is retrieved rapidly and revealed during automatic action selection
- Positional tuning of actions boosts the competitive gradient during planning

## Introduction

Producing a variety of movement sequences from memory fluently is an essential capacity of primates, in particular humans. It enables a skilled and flexible interaction with the world for a range of everyday activities - from tool-use, speech and gestural communication, to sports and music. Key to fluent sequence production is sequence planning before the initiation of the first movement [1,2], with longer preparation time benefitting sequence execution, i.e. reducing initiation time after a ‘Go’ cue and improving accuracy [3]. However, the underlying nature and content of sequence planning is still debated [4].

Computational models of sequence control, such as competitive queuing (CQ) models, suggest that preparatory activity reactivates sequence segments *concurrently* by means of a parallel activation gradient in the parallel planning layer [5]. Here the neural activation pattern for each sequence segment is weighted according to its temporal position in the sequence [6,7]. A rich literature indirectly supporting CQ in sequence control stems from observations of serial recall including transposition of neighbouring items and items occupying the same position in different chunks[6,8.9], and excitability of forthcoming items during sequence production[10]. Moreover, the CQ account has also been substantiated directly at the neurophysiological level in the context of well-trained finger sequences [11,12], saccades [13] and drawing geometrical shapes [14]. Importantly, these results have demonstrated that the neural gradient during planning is relevant to subsequent execution. In particular, response separation in the competitive gradient during sequence planning is predictive of sequence production accuracy [11,14]. Together, these data suggest that skilled sequence production involves the concurrent planning of several movements in advance before sequence initiation to achieve fluent performance.

While neural CQ during planning has been shown to predict subsequent production, it remains unclear which properties of the sequence this preparatory pattern encapsulates – the accuracy of the sequence (fluency of initiation and production quality), or the temporal structure of the sequence (speed and temporal grouping). Some CQ models assume the presence of a temporal context layer and that the activity gradients are learned by associations of the latter to each sequence element in the parallel planning layer, e.g. through Hebbian learning [7]. The form of activity in the context layer can be as simple as a decaying start signal [15], a combination of start and end signals [5,16] or a sequence of overlapping states [7,17]. Although primarily encoding serial order of sequence items, models utilizing overlapping states can implement effects of temporal grouping or sequence rhythm [7]. Therefore, it is possible that the competitive activation of actions during sequence planning encodes the temporal structure of the upcoming sequence.

In order to investigate the nature of sequence preparation and its relation to subsequent performance, we have developed a behavioural paradigm to capture the preparatory state of each item during planning of a well-learned sequence. Following training, participants prepared a motor sequence from memory following an abstract visual stimulus associated with a particular sequence of finger presses performed with a particular temporal structure and speed. In half of the trials during the test phase, the ‘Go’ cue was replaced by a finger press cue probing presses occurring at different positions of the sequence. We used reaction time (RT) and finger press accuracy to these ‘probes’ to compute as measures of the relative activation of planned actions during sequence planning.

We hypothesized that if competitive queuing primarily reflected the accuracy of the sequence plan, we would on average observe an enhancement of the CQ gradient with longer preparation time, as well as a correlation of the gradient with measures of sequence fluency and skill, specifically more rapid sequence initiation of correct sequences after the ‘Go’ cue, more accurate timing and fewer finger press errors. If, however, the gradient reflected the temporal structure – the speed and temporal grouping of the sequence, we should see that it is less pronounced for sequences twice as fast (speed manipulation), and shortened vs lengthened inter-press-intervals (IPI; temporal structure manipulation), because the actions are closer together in time.

We find that the relative level of activation of probed actions at the end of the planning period accords with their intended serial position. Contrary to the timing hypothesis, we found no reliable association with speed or temporal structure of the sequences. In contrast, we report that the corresponding CQ gradient is enhanced with longer preparation time, and is correlated with faster initiation of correct sequences and better temporal accuracy. Our data suggests that the competitive queuing gradient during planning primarily encodes the intended order of actions and the accuracy of a sequence plan, and not its overall speed or temporal structure. Based on this data, we propose a computational model that explains how the width of purely positional tuning could act on the relative activation state of actions during sequence planning independently of timing to enable accurate and fluent sequence performance.

## Results

### Finger press accuracy in sequences produced from memory was matched across conditions

Across three experiments, participants were trained for two days to associate two or three abstract visual cues with a particular four-element finger sequence performed with a particular temporal structure (Timing: slow, fast or irregular) following a brief preparation period (Delay between *Sequence* and *Go* cue onsets: short / 500ms, intermediate / 1000 ms and long / 1500 ms). In the test phase on the third day, they produced the respective sequences entirely from memory (Figure 1, all panels).

**Figure 1.**
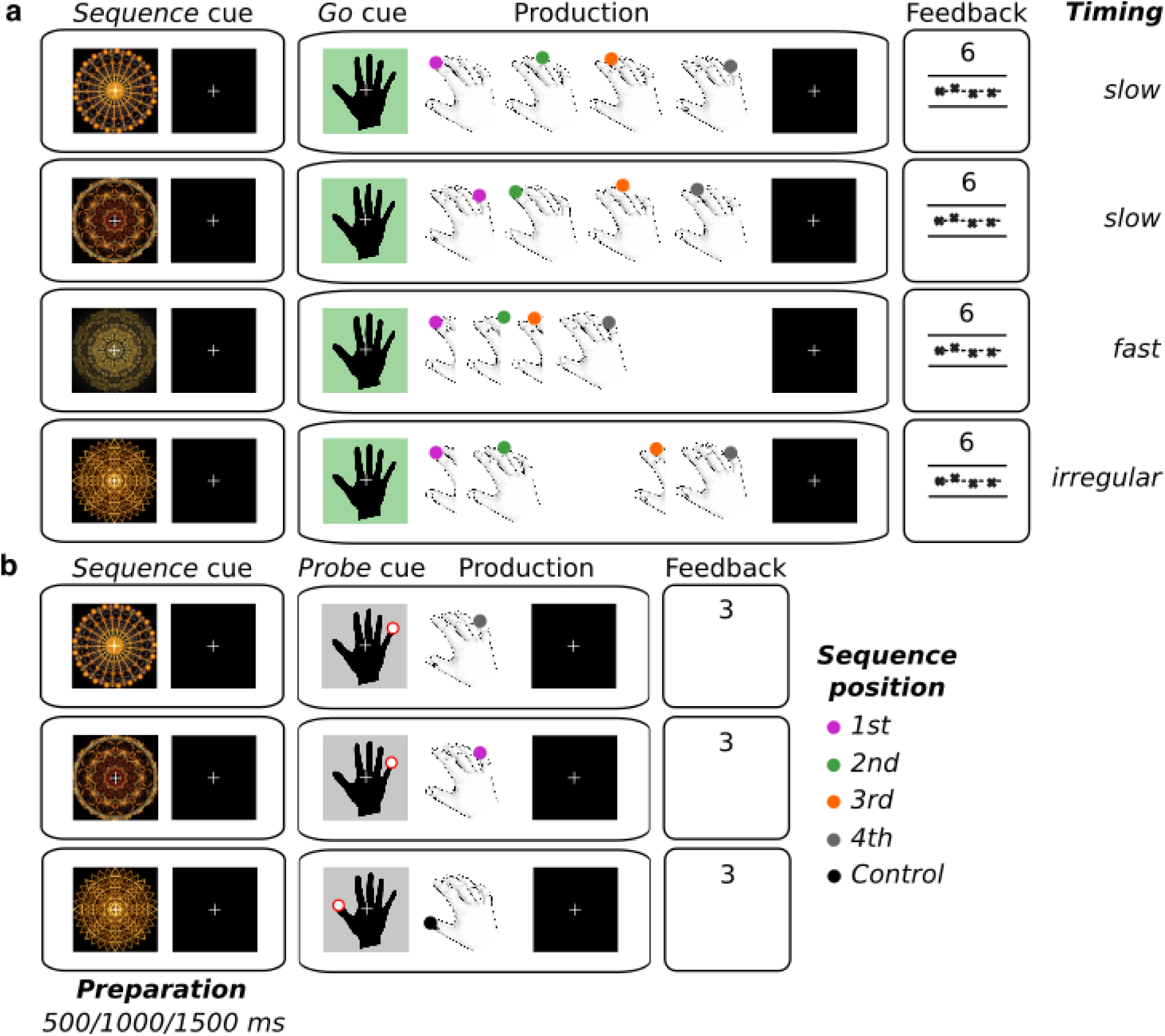
Experimental conditions. **a**. Participants were trained to produce 4-element finger sequences following a *Go* cue from memory. Each finger sequence or timing corresponded to a unique abstract visual *Sequence* cue presented up to 1500 ms before the *Go* cue (Preparation period). Experiment 1 cued the production of sequences with two different finger press orders. Here we manipulated the duration of the Preparation period (500, 1000 or 1500 ms). Experiments 2 and 3 had a fixed preparation duration of 1500 ms, but *Sequence* cues prompted the production of sequences with a different temporal structure (slow, fast and irregular). Participants received visual feedback in each trial on the accuracy of the press order and their timing, and received points based on press accuracy, temporal accuracy and initiation time (cf. Methods section). **b**. In all experiments, we introduced *Probe* trials, in which following the preparation period the *Go* cue was replaced with a *Probe* cue prompting a particular finger digit to be pressed, corresponding to each sequence position or control (thumb, which did not feature in any sequence production). This condition was used to obtain the reaction time (RT) and error rate for each position at the end of the preparation period. They received points for accurate production and fast RTs.

The finger error rate in sequence production from memory was higher in Experiment 1 than in Experiments 2 and 3. This is likely due to Experiment 1 involving the production of two different finger sequences produced with the same timing, and Experiments 2 and 3 involving the production of one finger sequence with different timings. The mean occurrence of finger errors, as indicated by either incorrect finger order or incomplete sequences, ranged from 0% to 26.6% in the short (*M* = 5.6%, *SD* = 6.9), 0% to 21.9% in the intermediate (*M* = 5.7% *SD* = 6), and from 0% to 15.6% in the long Delay condition (*M* = 4.6%, *SD* = 4.5) in Experiment 1. In Experiment 2, finger error rate varied between 0% and 5.5% at the slow timing (*M* = 2.2%, *SD* = 2.1), between 0% and 8.5% at the fast timing (*M* = 2.2%, *SD* = 2.5), and between 0% and 7% at the irregular timing (*M* = 2.3%, *SD* = 2.7). Error performance in Experiment 3 showed a rate between 0% and 13.3% at the slow timing (*M* = 2.6%, *SD* = 3.4), between 0% and 7.5% at the fast timing (*M* = 2.8%, *SD* = 2.5), and between 0% and 9.2% at the irregular timing (*M* = 2.4%, *SD* = 2.6).

Neither Delay (Experiment 1, *F* (2, 36) = .993, *p* = .451, *ηp*^*2*^ = .052) between the *Sequence* and the *Go* cue, nor the sequence Timing condition affected finger press accuracy during sequence production (Experiment 2, *F* (2, 34) = .006, p = .994, ηp^2^ = .000; Experiment 3, *F* (1.458, 24.787) = .249, *p* = .711, *ηp*^*2*^ = .014, Greenhouse-Geisser corrected, χ^2^ (2) = 7.436, *p* = .024, Figure 2c). This means that participants learned and prepared the finger order of all target sequences with the same finger accuracy, regardless of the preparation time or the temporal structure of the planned sequence.

**Figure 2.**
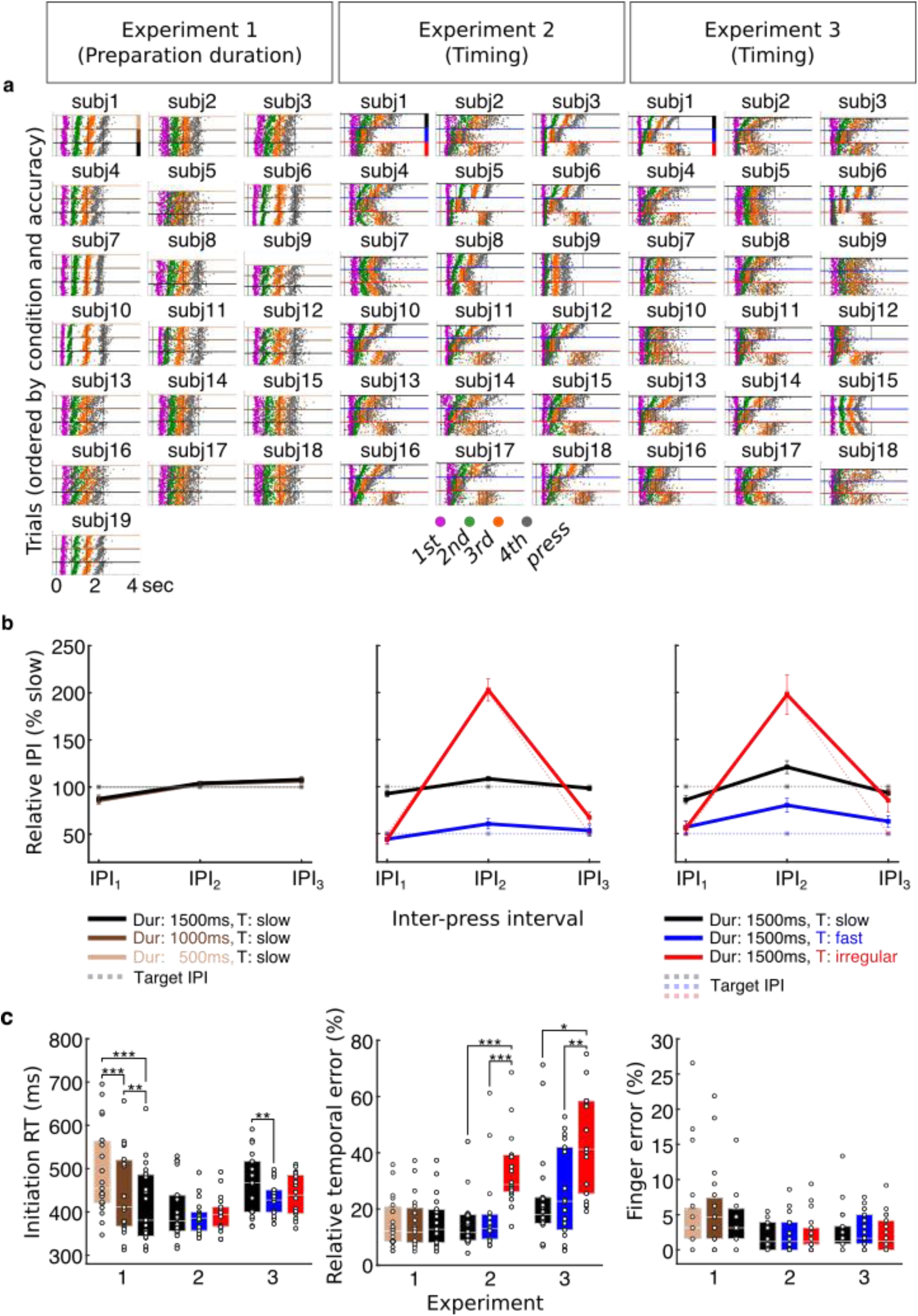
Sequence production. **a**. Individual participants’ raster plots show the timing of single button presses for each correct sequence trial produced from memory after the *Go* cue (t = 0) following training (target timing superimposed, grey lines). The colour code of the button presses corresponds to the press position in Figures 1 and 5. Within each condition, trials are ordered from most accurate to least accurate with regard to target onset (colour coding for conditions, cf. side bars in first participant panels, respectively). **b**. On average IPI production followed the target IPI structure, with a slow, twice as fast and an irregular sequence. IPIs were normalized across trials relative to slow isochronous condition. Preparation duration did not modulate IPI production of slow sequences, in contrast to the timing conditions. Error bars represent standard errors. **c**. Sequence initiation RT (*Go* cue to first press latency) decreased with preparation time (foreperiod effect). Relative temporal error was elevated for the irregular sequence in both experiments 2 and 3, in which it occurred. There was no effect of any of the conditions on finger press error rate, defined as proportion of incorrect trials. | * P ≤ 0.05 | ** P ≤ 0.01 | *** P ≤ 0.001.

### Participants produced sequences from memory with correct relative timing

When producing the sequences from memory during the test phase, participants had a general tendency to produce faster versions of the finger press sequences (Figure 2a), similar to the effects found in previous work [18]. The produced sequence duration was shorter than the target duration by 28.4% (SD = 10.7%), 38.6% (SD = 20.6%) and 46.2% (SD = 17.4%) in Experiment 1, 2 and 3, respectively (Figure 2a). However, the goal of our experimental design was to train participants to either retain or to modulate the *relative* timing across conditions according to the target relative IPIs, respectively (Figure 2b). Importantly, the majority of participants produced the sequences with the correct relative timing across conditions – on average the same temporal structure (slow) across preparation durations in Experiment1, and three different temporal structures (slow, fast, irregular) in Experiments 2 and 3 (Figure 2b).

Despite a largely overlapping sequence timing across preparation durations in Experiment 1 (Figure 2b, left), we found a small, but significant IPI × Delay interaction (3×3 repeated measures ANOVA: *F* (4, 72) = 2.528, *p* = .048, *ηp*^*2*^ = .123) explained by a modulation of 9 ms. Post-hoc comparisons (Bonferroni-corrected for nine tests) revealed a significant shortening of the 1^st^ interval in the short (*p* = .002) and in the intermediate delay (*p* = .002) compared to the long delay conditions. Additionally, the 3^rd^ interval was larger in the long delay than at the intermediate delay (*p* = .004). This shows that there was a tendency to slightly compress the 1^st^ interval with shorter preparation time and slightly expand the 3^rd^ interval with longer preparation time. However, the size of this temporal modulation in Experiment 1 was 97% smaller than the temporal structure modulation induced in Experiments 2 and 3. Any potential sequence timing effect on CQ activation of actions during preparation should thus be vastly augmented in Experiments 2 and 3.

As expected, Experiment 2 showed a significant IPI × Timing interaction (3×3 repeated measures ANOVA: *F* (1.260, 21.417) = 59.485, *p* < .001, *ηp*^*2*^ = .778, Greenhouse-Geisser corrected, χ^2^ (9) = 97.832, *p* < .001). The pairwise comparisons (Bonferroni-corrected for nine tests) of the produced intervals confirmed that the participants modulated their relative interval production according to the trained target interval structure. In accordance with the target sequence, in the slow timing condition the 1^st^ interval was significantly longer than in the fast (*p* < .001) and the irregular timing conditions (*p* < .001), while there was no difference between the fast and irregular conditions (*p* = 1.000) for the latter. The 2^nd^ interval length increased slightly, yet proportionally for both the slow and the fast conditions, retaining the significant difference (*p* < .001) and doubled in length for irregular relative to the slow timing condition (*p* < .001). Finally, the 3^rd^ interval exhibited a very similar profile to the 1^st^ interval (slow *vs* fast, *p* < .001; slow *vs* irregular, *p* < .001), but showed a slightly lower percent interval for the fast compared to the irregular conditions (*p* = .027). Overall, the IPI production data shows that the fast sequence was on average half as long as the slow sequence, and the irregular timing condition changed the relative interval structure from regular slow to an irregular short–long–short pattern of the same sequence duration. Experiment 3 replicated the findings of Experiment 2 showing a significant IPI × Timing interaction (3×3 repeated measures ANOVA: *F* (1.558, 26.485) = 17.369, *p* < .001, *ηp*^*2*^ = .505, Greenhouse-Geisser corrected, χ^2^ (9) = 61.311, *p* < .001). Again, post-hoc pairwise comparisons (Bonferroni-corrected for nine tests) confirmed that the 1^st^ interval of the slow condition was longer than that of the fast (*p* = .001) and irregular (*p* = .003) conditions, while no difference was found between fast and irregular conditions (*p* = 1.000). The 2^nd^ interval was significantly longer in the slow compared to the fast condition (*p* = .001), but shorter compared to the irregular condition (*p* = .005). Similarly, the fast condition was half as long in the irregular condition (*p* < .001). The 3^rd^ interval was twice as long for the slow relative to the fast condition (*p* < .001), but failed to show a significant shortening for the irregular relative to the regular slow sequence conditions (*p* = 1.000), and there was only a marginally significant difference between fast and irregular conditions (*p* = .096). Overall, our findings demonstrate that, on average, participants retrieved and produced the finger sequences form memory with distinct temporal structures according to the relative timing of slow, fast and irregular target intervals.

### Longer preparation duration speeds up sequence initiation

The time to initiate a correct action sequence after the *Go* cue can be taken as a marker of the state of action planning after the preparatory delay [3,19.20]. We found a significant difference in mean initiation RT with Delay (Experiment 1, one-way repeated measures ANOVA: *F* (1.382, 24.877) = 52.809, *p* < .001, *ηp*^*2*^ = .746, Greenhouse-Geisser corrected, χ^2^ (2) = 10.074, *p* = .006) (Figure 2c, left). Pairwise comparisons (Bonferroni-corrected for three tests) confirmed that initiation time for the intermediate and long delay conditions was significantly shorter than following the short delay (intermediate *vs* short delay, *p* < .001; long *vs* short delay, *p* < .001). Similarly, sequence initiation following a long delay performed at significantly faster mean RT as compared to the intermediate delay period (*p* = .005). Notably, this effect is also in line with the classic foreperiod effect identified for single actions showing faster RTs for longer foreperiod durations [21,22] suggesting that temporal expectation of the *Go* cue may also contribute to faster sequence initiation in addition to the state of sequence planning.

In contrast to the effect of preparation time, the planned temporal structure of the sequence did not consistently affect sequence initiation RT (Figure 2c, left). There was no main effect of sequence Timing in Experiment 2 (one-way repeated measures ANOVA: *F* (1.407, 23.917) = 1.700, *p* = .207, *ηp*^*2*^ = .091, Greenhouse-Geisser corrected, χ^2^ (2) = 8.759, *p* = .013), but a main effect of Timing in Experiment 3 (one-way repeated measures ANOVA: *F* (1.294, 21.993) = 11.590, *p* = .001, *ηp*^*2*^ = .405, Greenhouse-Geisser corrected, χ^2^ (2) = 12.632, *p* = .002). Specifically, as explained by pairwise comparisons (Bonferroni-corrected for three tests), participants in Experiment 3 were slower at initiating a slow regular sequence when compared to a fast regular sequence (*p* = .006) and an irregular sequence (*p* = .010) whilst there was no difference in initiation RT between the fast regular and the irregular conditions (*p* = .118).

### Sequences involving irregular inter-press-intervals were produced with less accurate timing

Next, we aimed to establish whether preparation time and the temporal interval structure of sequences modulated the observed relative timing accuracy across presses (Figure 2c, middle). In Experiment 1, the mean relative temporal error did not differ across the three delay conditions (one-way repeated measures ANOVA: *F* (2, 36) = .105, *p* = .901, *ηp*^*2*^ = .006). This indicates that time to prepare the sequence did not affect the degree of relative temporal accuracy. Here, sequence accuracy in conditions with a shorter preparation time might have been compensated by slower initiation RT (cf. above). In Experiment 2, there was a significant effect of Timing (one-way repeated measures ANOVA: *F* (2, 34) = 28.226, *p* < .001, *ηp*^*2*^ = .624). Pairwise comparisons (Bonferroni-corrected for three tests) revealed that participants performed at a higher relative temporal accuracy when producing a slow regular sequence compared to an irregular (*p* < .001) and a fast regular compared to an irregular sequence (*p* < .001), while there was no difference between the slow and fast regular conditions (*p* = 1.000). Experiment 3 replicated the main effect of Timing (one-way repeated measures ANOVA: *F* (1.454, 24.723) = 7.060, *p* = .007, *ηp*^*2*^ = .293, Greenhouse-Geisser corrected, χ^2^ (2) = 7.527, *p* = .023). In line with the findings of Experiment 2, there were smaller temporal errors in the slow sequence condition compared to the irregular condition (*p* = .049) and in the fast compared to the irregular sequence (*p* =. 008). There was no significant difference in temporal performance between the two regular conditions (*p* = 1.000). These results suggest that the production of sequences which consist of several different IPIs (irregular sequence) as opposed to only one interval length (isochronous/regular sequence) is associated with decreases in temporal accuracy of the sequence.

### Action probes show graded activation of sequence elements at the end of preparation and are modulated by preparation duration, not sequence timing

In half of the trials in the test phase on Day 3, instead of the *Go* cue that prompted the production of the prepared finger sequence, participants encountered a visual *Probe* cue which instructed them to respond with the corresponding finger press as quickly and accurately as possible (Figure 1b). This allowed us to obtain two behavioural measures related to the competitive state of each constituent press at the end of the preparation period – the relative action activation and the probability of correct action selection – for each action in the sequence, respectively. Specifically, lower RT would suggest higher activation of the correctly selected action and lower press error rate a higher probability of selection, and thus availability of the associated action.

#### Reaction times

To evaluate the relative activation state of the probed actions associated with different sequence positions (Figure 3a), we normalized the RT of each probe position relative to the RT of the first position of the prepared sequence (RT increase in % relative to 1^st^ position; cf. Supplementary Figure 1a for raw RT values). In Experiment 1, results showed a main effect of Position (*F* (1,615, 29.062) = 45.958, *p* < .001, *ηp*^*2*^ = .719, Greenhouse-Geisser corrected, χ^2^ (5) = 22.621, *p* < .001). Planned contrasts to detect differences between adjacent positions revealed a significant RT increase with each position compared to its preceding one (2^nd^ *vs* 1^st^ position, *F* (1, 18) = 63.360, *p* < .001, *ηp*^*2*^ = .779; 3^rd^ *vs* 2^nd^ position, *F* (1, 18) = 54.534, *p* < .001, *ηp*^*2*^ = .752; 4^th^ *vs* 3^rd^ position, *F* (1, 18) = 24.900, *p* < .001, *ηp*^*2*^ = .580). In line with our hypothesis, these differences indicate a graded activation of actions according to their serial position in the sequence, with the first action being the most activated. We also found a significant Position × Delay interaction (*F* (6, 108) = 2.980, *p* = .010, *ηp*^*2*^ = .142). Planned contrasts using the long preparation duration as the reference condition for the Delay factor, showed a significantly greater increase of the 2^nd^ position relative to the 1^st^ position at the long *vs* the short delay (*F* (1, 18) = 7.349, *p* = .014, *ηp*^*2*^ = .290). These results suggest that a longer preparation time prior to sequence execution boosted the competitive activation of actions.

**Figure 3.**
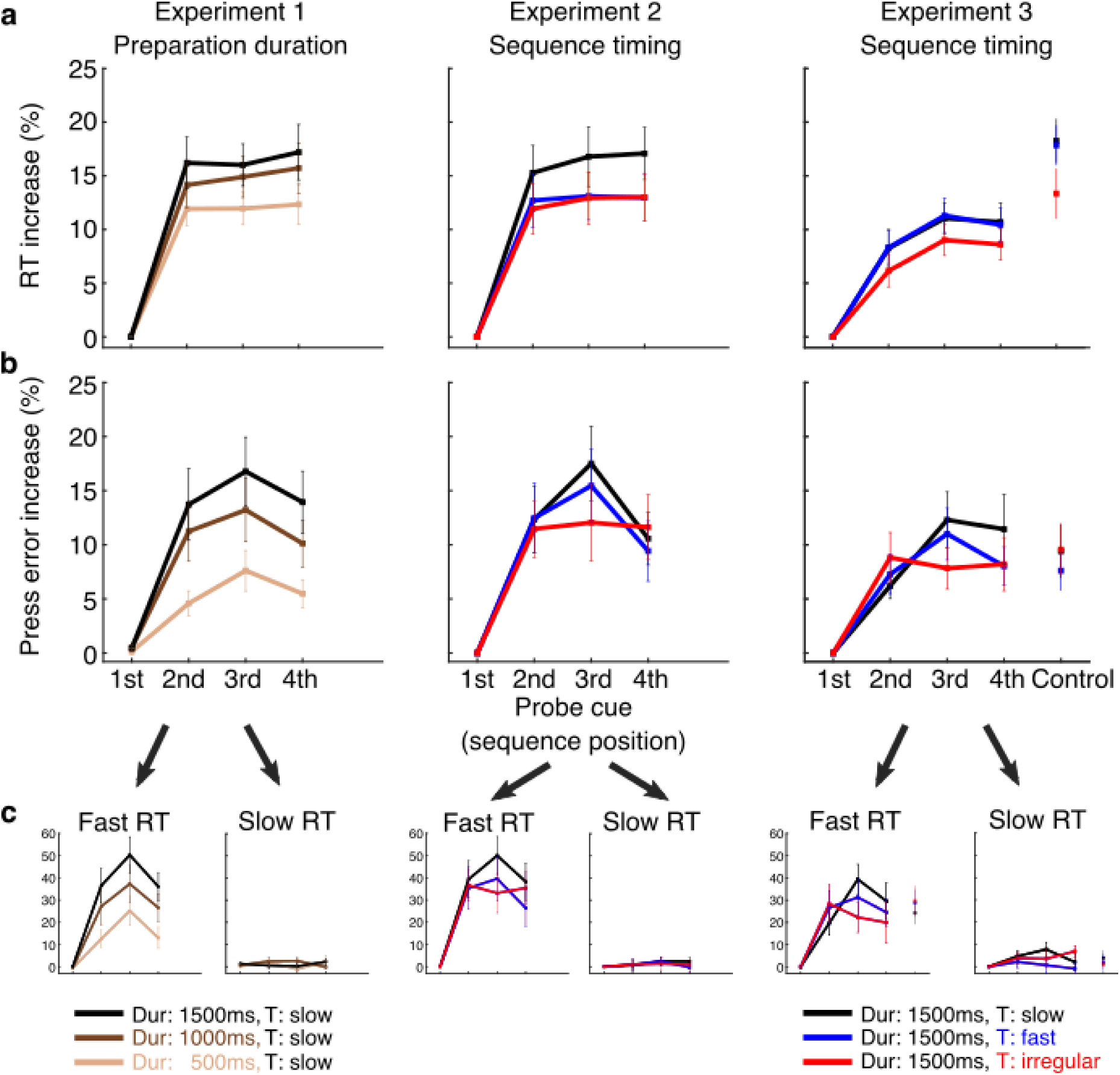
Competitive state of actions during sequence preparation. *Probe* trials prompting the production of an action associated with the 1^st^-4^th^ press position of the prepared sequence or a control action not present in any sequence (Experiment 3). **a**. Reaction time (RT) gradually increased for later sequence positions relative to the first position and became more pronounced with longer preparation duration, with responses to actions in later sequence positions becoming slower on average, when participants had more time to prepare the sequence (Experiment 1). No significant changes to RT increase were observed between conditions in which participants prepared sequences performed with different timing (Experiment 2 and 3). The RT increase was most pronounced for the action not featuring in the planned sequence (control action was a thumb press; Experiment 3), cf. raw RT graphs in Supplementary Figure 1. **b**. Press error rate also increased gradually for later sequence positions relative to the first position, with the exception of the last position, thus approaching an inverted U-shape. The press error gradient became more pronounced with longer preparation duration, i.e. responses to probes associated with later positions became less accurate when participants had more time to prepare the cued sequence. **c**. This pronounced effect on error increase was driven by trials where the response to action probes was short (lower RT quartile). When participants slowed down their response allowing more time for deliberation and correction (upper RT quartile), the characteristic error increase was absent or less pronounced. Error bars represent standard errors.

In Experiment 2, we replicated the main effect of Position (*F* (2.230, 37.904) = 25.131, *p* < .001, *ηp*^*2*^ = .596, Greenhouse-Geisser corrected, χ^2^ (5) = 15.333, *p* = .009). Planned contrasts showed significant differences when contrasting all pairs of adjacent probe positions replicating the findings of Experiment 1. Specifically, RT increase of the 2^nd^ position was larger compared to the 1^st^ (*F* (1, 17) = 48.072, *p* < .001, *ηp*^*2*^ = .739). Similarly, there was a significantly greater RT increase for the 3^rd^ position *vs* the 2^nd^ (*F* (1, 17) = 32.040, *p* = .001, *ηp*^*2*^ = .653) and the 4^th^ *vs* the 3^rd^ (*F* (1, 17) = 28.873, *p* < .001, *ηp*^*2*^ = .629). Crucially, there was no interaction between Position and Timing (*F* (2.430, 41.318) = 2.823, *p* = .061, *ηp*^*2*^ = .142, Greenhouse-Geisser corrected, χ^2^ (20) = 59.308, *p* < .001). This suggests that preparing a sequence with a different temporal structure did not impact the competitively cued activations at the end of preparation.

Experiment 3 once more replicated a main effect of Position (*F* (3, 51) = 29.852, *p* < .001, *ηp*^*2*^ = .637). Planned contrasts, similarly, revealed a significantly greater RT increase for the 2^nd^, 3^rd^ and 4^th^ positions over their preceding 1^st^, 2^nd^ and 3^rd^ positions, respectively (2^nd^ *vs* 1^st^, *F* (1, 17) = 61.485, *p* < .001, *ηp*^*2*^ = .783; 3^rd^ *vs* 2^nd^, *F* (1, 17) = 69.762, *p* < .001, *ηp*^*2*^ = .804; *F* (1, 17) = 14.180, *p* = .002, *ηp*^*2*^ = .455). In addition, the finger press which did not feature in any of the planned sequences (control action: thumb) showed a further RT increase relative to the last (4^th^ position) item of the sequence (paired samples t-test: *t* (17) = 3.062, *p* = .007, *d* = .840, two-tailed). Timing did not interact with Position (*F* (3.743, 63.632) = 1.089, *p* = .367, *ηp*^*2*^ = .060, Greenhouse-Geisser corrected, χ^2^ (20) = 36.727, *p* = .014). This result replicates Experiment 2. Thus, CQ during sequence preparation was not dependent upon the speed or temporal structure of the planned sequences, and suggests that fine grained competitive activation gradient of constituent actions of a sequence is activated above the level of an unrelated effector.

#### Error rate

To evaluate the relative probability of selection of the probed actions associated with different sequence positions (Figure 3b), in each experiment the finger error rate for each probe action was calculated and normalized to that of the first position (Error rate increase in % relative to 1^st^ position; cf. Supplementary Figure 1b for raw error rate values). Equally, we assessed the same factors, predicting an ascending error rate increase by position. Experiment 1 showed a main effect of Position (4 × 3 repeated measures ANOVA: *F* (1,948, 35.073) = 18.017, *p* < .001, *ηp*^*2*^ = .500, Greenhouse-Geisser corrected, χ^2^ (5) = 13.595, *p* = .019). Planned contrasts revealed a significantly increased error rate for the 2^nd^ position compared to the 1^st^ position (*F* (1, 18) = 29.675, *p* < .001, *ηp*^*2*^ = .622) and for the 3^rd^ position compared to the 2^nd^ position (*F* (1, 18) = 25.937, *p* < .001, *ηp*^*2*^ = .590), whilst the last, 4^th^ position showed a lower error increase than the 3^rd^ position (*F* (1, 18) = 5.092, *p* = .037, *ηp*^*2*^ = .220). This error rate pattern during preparation shows an inverted U-shape, similar to serial position curves during production [23] and suggests the presence of a ranked probability across sequence positions for action selection.

The Position × Delay interaction (*F* (4.137, 74.466) = 3.813, *p* = .007, *ηp*^*2*^ = .175, Greenhouse-Geisser corrected, χ^2^ (20) = 34.036, *p* = .028) was driven by a significant increase of the 2^nd^ position compared to 1^st^ position at the long *vs* the short delay (*F* (1, 18) = 10.877, *p* = .004, *ηp*^*2*^ = .377) as revealed by planned contrasts. No other pairs showed a significant difference. In accordance with the RT results, these findings suggest that accuracy of probe elements during sequence planning is modulated by preparation, in that a longer preparation time is associated with more pronounced error increases for the 2^nd^ position when compared to a short preparation period, suggesting less availability for selection of actions in later positions the more sequence planning advances.

Experiment 2 showed a significant main effect of Position (*F* (3, 51) = 14.397, *p* < .001, *ηp*^*2*^ = .459). Planned contrasts revealed that this effect was driven by a significant error rate increase for the 2^nd^ position *vs* the 1^st^ position (*F* (1, 17) = 24.070, *p* < .001, *ηp*^*2*^ = .586). The 3^rd^ position performed at a greater error increase than the 2^nd^ position (*F* (1, 17) = 15.510, *p* = .001, *ηp*^*2*^ = .477), whilst the 4^th^ position was not significantly different from the 3^rd^ position (*F* (1, 17) = 1.284, *p* = .273, *ηp*^*2*^ = .070). These results replicate the CQ effect of serial actions during preparation, found in Experiment 1, with a graded increase in error rates for later elements up to the 3^rd^ position. We did not find a significant Position × Timing interaction (*F* (6, 102) = 1.583, *p* = .160, *ηp*^*2*^ = .085).

Similarly in Experiment 3, there was a main effect of Position (*F* (3, 51) = 13.725, *p* < .001, *ηp*^*2*^ = .447), with the 2^nd^ position showing a greater error increase than the 1^st^ position (*F* (1, 17) = 29.074, *p* < .001, *ηp*^*2*^ = .631), and the 3^rd^ position performing with more errors than the 2^nd^ position (*F* (1, 17) = 17.903, *p* = .001, *ηp*^*2*^ = .513). Error rates of the 4^th^ position did not differ from the 3^rd^ position (*F* (1, 17) = 3.791, *p* = .068, *ηp*^*2*^ = .182). Our prediction that the control action would not be part of this queuing pattern, implying a much weaker probability to be selected for execution, was refuted by a non-significant difference from the 4^th^ position (paired samples t-test: *t* (17) = -.323, *p* = .751, *d* = .111, two-tailed). As in Experiment 2, Position did not interact with Timing (*F* (3.803, 64.654) = 1.869, *p* = .130, *ηp*^*2*^ = .099, Greenhouse-Geisser corrected, χ^2^ (20) = 42.899, *p* = .002). Together, this indicates that the competitive error rate for probed actions during preparation was not modulated by the speed or temporal structure of the planned sequence.

Whilst there was no significant interaction of Timing and Position for action probes during preparation (neither for RT, nor for error rate), we observed a non-significant, but consistent flattening of the CQ curve for the temporally irregular sequence across Experiments 2 and 3. However, this patterns of results cannot be attributed to changes in temporal grouping of actions per se, but may be driven by accuracy: The irregularly (non-isochronously) timed sequence was characterized by a highly significant increase in relative temporal error when compared to both the slow and fast regularly (isochronously) timed sequences (Figure 2b, Experiments 2 and 3) due to the increased temporal complexity. This lends support to the alternative hypothesis, namely that the precision of the sequence plan is driving the CQ state of actions during the preparation period.

### A steeper CQ error gradient is bound to fast responses

Next, we sought to determine whether the characteristic error rate gradients were the result of automatic responses, or deliberated action selection after the *Probe* cue. To test this hypothesis, we assessed the error rate gradient for fast *vs* slow responses following the *Probe* cue. We extracted the relative error rate increases for action probes in the first and third RT quartiles for each experiment (Figure 3c). Only for the fast responses, we found a main effect of Position (Experiment 1, *F* (1.758, 31.650) = 19.731, *p* < .001, *ηp*^*2*^ = .523, Greenhouse-Geisser corrected, χ^2^ (5) = 17.279, *p* = .004; Experiment 2, *F* (2.033, 34.559) = 16.325, *p* < .001, *ηp*^*2*^ = .490, Greenhouse-Geisser corrected, χ^2^ (5) = 19.928, *p* = .001; Experiment 3, *F* (3, 51) = 12.749, *p* < .001, *ηp*^*2*^ = .429). Planned contrasts for adjacent positions confirmed a graded increase in finger errors up to the 3^rd^ position (Experiment 1, 2^nd^ position *vs* 1^st^ position, *F* (1, 18) = 26.954, *p* < .001, *ηp*^*2*^ = .600; 3^rd^ position *vs* 2^nd^ position, *F* (1, 18) = 35.745, *p* < .001, *ηp*^*2*^ = .665; 4^th^ position *vs* 3^rd^ position, *F* (1, 18) = 2.347, *p* = .143, *ηp*^*2*^ = .115; Experiment 2, 2^nd^ position *vs* 1^st^ position, *F* (1, 17) = 27.138, *p* < .001, *ηp*^*2*^ = .615; 3^rd^ position *vs* 2^nd^ position, *F* (1, 17) = 15.222, *p* = .001, *ηp*^*2*^ = .472; 4^th^ position *vs* 3^rd^ position, *F* (1, 17) = 4.982, *p* = .039, *ηp*^*2*^ = .227; Experiment 3, 2^nd^ position *vs* 1^st^ position, *F* (1, 17) = 21.580, *p* < .001, *ηp*^*2*^ = .559; 3^rd^ position *vs* 2^nd^ position, *F* (1, 17) = 13.888, *p* = .002, *ηp*^*2*^ = .450; 4^th^ position *vs* 3^rd^ position, *F* (1, 17) = 1.845, *p* = .192, *ηp*^*2*^ = .098). We also found a significant Position × Delay interaction (Experiment 1, *F* (6, 108) = 4.003, *p* = .001, *ηp*^*2*^ = .182), driven by a significant increase of the 2^nd^ position 1^st^ position at the long *vs* the short delay (*F* (1, 18) = 18.132, *p* < .001, *ηp*^*2*^ = .502) and at the long *vs* the intermediate delay (*F* (1, 18) = 10.370, *p* = .005, *ηp*^*2*^ = .366). Error rate increases did not change by sequence position number in the slow responses (Experiment 1, *F* (3, 54) = .313, *p* = .816, *ηp*^*2*^ = .017; Experiment 2, *F* (3, 51) = .552, *p* = .649, *ηp*^*2*^ = .031; Experiment 3, *F* (3, 51) = 1.672, *p* = .185, *ηp*^*2*^ = .090), accompanied by an absent Position × Delay interaction (Experiment 1, *F* (3.559, 64.058) = 1.302, *p* = .280, *ηp*^*2*^ = .067, Greenhouse-Geisser corrected, χ^2^ (20) = 37.340, *p =* .012). The control action was not different from the 4^th^ position in either RT pole (fast RTs, *t* (17) = 1.654, *p* = .117, *d* = .473, two-tailed; slow RTs, *t* (17) = .203, *p* = .842, *d* = .050, two-tailed). These results suggest that the CQ gradient at the end of a preparation period of 500 to 1500 ms was driven by automatic responses rather than by cognitive action selection and replanning, and constitute a readout for the state of actions during sequence planning.

### Preparatory CQ gradient correlates with temporal accuracy and initiation speed

Neurally derived CQ of sequence actions during planning predicts the participants’ subsequent performance accuracy as shown previously[11]. In line with this finding, we found that more time to prepare a sequence is associated with a more pronounced competitive RT and error rate increase for action probes. To test the association directly, we predicted that a more pronounced (steeper) CQ gradient of RTs and error rates would correlate with a better performance in sequence production, specifically with faster correct sequence initiation, and less temporal and finger press errors. Correlation analyses were performed on group data obtained from trials in the slow timing condition and long preparation duration (1500 ms) present across all three experiments (N = 55). The magnitude of the CQ gradient during preparation was calculated based on RT and error rate increase data in *Probe* trials (difference between adjacent positions; Figure 4a and 4b). Results showed that participants with a more pronounced CQ based on relative RT and error rate increase initiated correct sequences faster (CQ RT increase: *r* = -.393, *p* = .002, one-tailed; CQ error increase: *r* = -.539, *p* < .001, one-tailed). Higher RT based CQ also predicted smaller relative temporal error (*r* = -.345, *p* = .005, one-tailed). However, in contrast to the correlations with the neural measure of CQ reported in a previous study[11], neither CQ RT increase, nor CQ error increase showed negative correlations with finger error (CQ RT increase: *r* = .083, *p* = .273, one-tailed; CQ error increase: *r* = .118, p = .196, one-tailed). Also, contrary to the CQ RT increase, a more pronounced CQ error increase of probe actions at the end of preparation was not associated with reduced temporal error during execution (*r* = -.051, p = .356, one-tailed). Thus, CQ error increase may be a less sensitive predictor for temporal accuracy than CQ RT increase.

**Figure 4.**
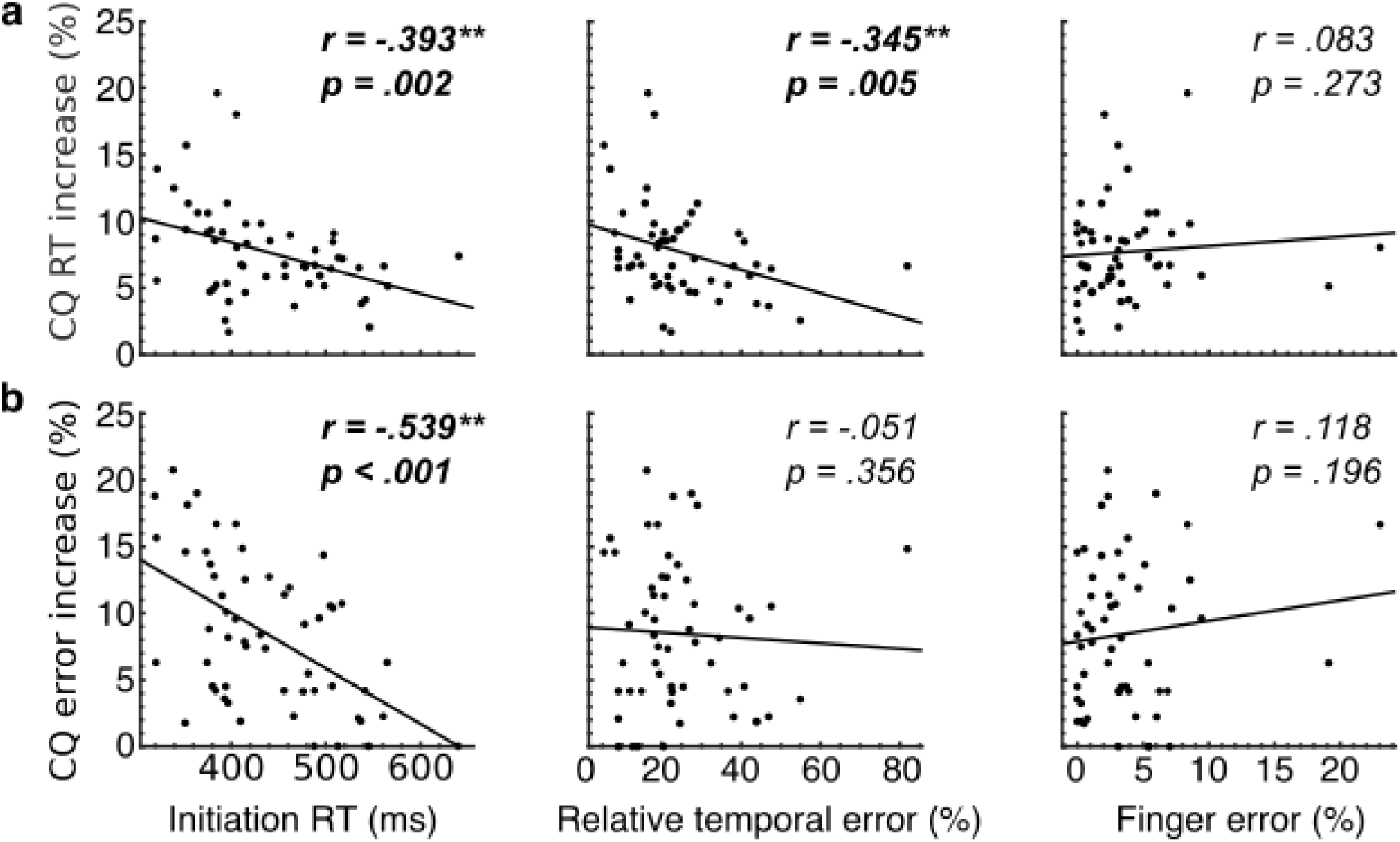
Correlation between overall CQ measures during preparation and subsequent production. Average relative RT (**a**) and press error increase (**b**) between adjacent positions (1^st^ to 2^nd^, 2^nd^ to 3^rd^, 3^rd^ to 4^th^) obtained through probe trials was taken as a proxy for CQ of actions during preparation. Larger CQ during preparation was associated with faster initiation speed of correct sequences, and smaller relative temporal errors. Larger CQ was not associated with reduced finger error rate (proportion of trials with wrong finger order, finger order repetitions, or missing presses), as predicted based on neural CQ findings of Kornysheva et al. [11]. All correlations are one-tailed, in line with one-sided predictions.

In sum, consistent with previous neurophysiological findings [11], our behavioural results show that during sequence preparation of sequence from memory participants establish a competitive activation and selection gradient of constituent actions according to their serial order. This competitive gradient expands with longer preparation durations and is more pronounced in participants with faster sequence initiation and more precise interval timing.

## Discussion

Sequence planning is central to skilled action control, however its content and organisation is poorly understood[4,24]. Neurophysiological findings in humans have demonstrated that a trained action sequence is pre-planned by establishing a competitive activation gradient of action patterns according to their serial position, and that the quality of this neural pattern during planning predicts subsequent performance [11–14]. Here we have established a behavioural measure of the preparatory action activation gradient and demonstrate that it reflects the skill (sequence production accuracy and fluency of initiation), rather than the temporal structure (sequence production speed and temporal interval pattern) of the planned sequence. Both the time to respond and the probability of making a finger press mistake increased progressively when participants responded to action cues during preparation that were associated with later *vs* earlier positions in the respective sequence. The non-linear increase was particularly pronounced for the first three out of four planned actions in the sequence. This response gradient demonstrates that the relative availability of each planned action in the sequence decreases with serial position, as predicted by competitive queuing (CQ) models [6,7,25,26] and previous neurophysiological findings [11,14].

The preparatory action activation gradient markedly contrasts with mechanisms for non-sequential action planning involving multiple actions: A cued set of possible actions triggers equal activity increase in cortical populations tuned to the respective actions, and the preparatory competition is only resolved once an action cue specifies the target action [27]. In contrast, sequence preparation establishes a fine-tuned gradient of action activations, with the latter switching flexibly depending on the retrieved sequence. Notably, actions that were part of the planned sequence were activated above the level of a control action which was not part of the retrieved sequence (Figure 3a, right). This suggests that all constituent actions were concurrently activated above a passive baseline, albeit to a different degree depending on their position in the planned sequence.

Our study provides a behavioural measure of the competitive state of constituent actions during sequence planning. This is complementary to previous behavioural work which revealed CQ of actions during production, such as accuracy and RT curves obtained from sequence execution [28,29], or on-the-fly action planning following sequence initiation, assessed behaviourally[30] and through measures of cortico-spinal excitability [10]. Gilbert and colleagues have employed a paradigm at the interface between sequence preparation and production to characterize the CQ profiles the respective sequential actions – silent rehearsal [31]. Here participants were asked to listen to sequences of spoken digits and silently rehearse the items during a retention interval. They received explicit instructions to rehearse the sequence at the same pace as active production. After an unpredictable delay, a tone prompted the report of an item being rehearsed at that moment and revealed graded overlapping probabilities of neighbouring items, suggesting potential CQ during internal rehearsal. In contrast to the latter study, our paradigm did not allow for active rehearsal during preparation: First, our participants retrieved the sequence entirely from memory without a sensory instruction period which might have facilitated active entrainment with the sequence prior to planning. Second, the period for sequence retrieval and planning was comparatively brief (ranging from 500 to 1500 ms after *Sequence* cue onset) and not sufficient to cycle through the full sequence at the rate participants employed for active production. In addition, if the observed CQ gradient were somehow driven by silent rehearsal at the target rate, it would have been more pronounced for the fast sequences, as more of the planned sequence could fit into the preparation phase. However, there was no significant difference between relative activation curves for fast and slow sequences.

Whilst active motor rehearsal at scale during the short preparation phase is unlikely, an alternative serial mechanism underlying the different levels of action activation may be mediated by rapid sequence replay. The latter has been observed in the hippocampus during navigation tasks [32] and perceptual sequence encoding [33], as well as in the motor cortex in the context of motor sequence learning tasks [34]. Replay has been shown to involve fast sweeps through the neural patterns associated with the sequence during wakeful rest and planning (preplay) [32,35–37], and is characterized by a multifold temporal sequence compression [33,34,38,39]. How replay could translate into a parallel activation of serial items described here is uncertain. One possibility is that serial sweeps during motor sequence preparation involve fast repeated replay fragments [38,40] of different length during planning, starting with the first elements – e.g. 1^st^-2^nd^-3^rd^, 1^st^-2^nd^, 1^st^, 1^st^-2^nd^-3^rd^-4^th^, 1^st^-2^nd^ etc. This would produce an overall bias towards the activation of earlier rather than later parts of the planned sequence, which may be translated into a cumulative ramping activity for each constituent action by a separate neuronal mechanism during the preparation period [27,41]. Future analysis of the ‘sequenceness’ [33,34] of the corresponding neural patterns during preparation should shed light on the presence of preplay and its hypothesized relationship to parallel CQ of actions [11].

The CQ gradient was established after a brief retrieval and preparation period, and revealed through faster rather than slower responses to probes (Figure 3c). This suggests that out behavioural measure of CQ during sequence planning reflects a rapid and automatic process involved in the execution of well-trained motor sequences from memory, and is not a result of slow deliberation or higher-level decision making. Contrary to a prominent account of motor control of skill learning [42,43], this data implies that discrete motor sequence production incorporates automatic planning mechanisms which are associated with fluent and accurate execution of sequential actions.

Remarkably, longer preparation times reinforced the competitive activation gradient making responses to action probes for later sequence positions even slower and more inaccurate relative to those for earlier positions. Whilst counterintuitive in the context of single action performance gains from longer foreperiod durations [21], the gradient expansion with time suggests a dynamic refinement of the plan for sequence production during the retrieval and preparation phase. This refinement involves the graded suppression of later actions in the sequence, making them less available for production, even more so with time. Crucially, the gradient increase with preparation duration was not accompanied by substantial expansion or compression of sequence production. Instead, the CQ gradient was associated with a faster initiation of correctly performed sequences whilst retaining the same level of press and timing accuracy, suggesting increased sequence fluency. This demonstrates that the action activation gradient established during planning reflects the preparedness for correct and fluent production, rather than the planned temporal structure of the sequence.

Furthermore, participants who had a more pronounced competitive activation during planning exhibited both faster initiation times and a more accurate temporal execution of the sequence after the “Go” cue, particularly when looking at the RT based CQ gradient. These findings strengthen the interpretation that an ordered competitive activation of actions during planning preempts subsequent fluency and temporal accuracy of the sequence [11]. Yet, we did not replicate the association of the planning gradient with finger error probability found in the latter study. This may be due to a smaller pool of timing and finger order sequences that the participants had to learn relative to the previous paradigm, and the presence of only one finger order (but different sequence timing) in Experiments 2 and 3. This likely facilitated finger accuracy to reach ceiling levels in a substantial number of participants. Future experiments should resolve an association with finger accuracy through the inclusion of a larger pool of trained sequences to provoke more frequent finger errors. Alternatively, reaching or drawing tasks would allow to make the spatial in addition to temporal feature of the sequential behaviour continuous and capture fine-grained spatial errors at overall high accuracy levels of sequence production.

In contrast, doubling the speed of sequence production did not change the relative activation between sequential actions at the end of the preparation period. This suggests invariance of the gradient in the competitive planning layer across sequences produced at different time scales. This transfer across speed profiles is in line with the presence of flexible motor timing and temporal scaling in dynamic neuronal populations [44,45], and a separate neural process controlling the speed of an action or action sequence during execution, e.g. through the strength of an external input to the network involved in the generation of timed behavior [44]. Preparing a sequence of the same length with an irregular compared to isochronous interval structure was associated with a tendency for a dampened CQ gradient during sequence planning. However, this non-significant trend on CQ is unlikely to be the effect of temporal grouping, as the irregular interval sequence was characterized by a significant increase in temporal interval production error (Figure 2c, middle panel). Instead, we hypothesize that longer preparation time (above 1500 ms) would have benefitted the participants and enhanced the relative activation gradient in line with Experiment 1 in order to form a more accurate plan for the complex sequencing of two different (non-isochronous) rather than just one constituent IPI (isochronous).

Our results show that CQ of actions during sequence planning reflects the overall action order and temporal accuracy of the sequence, but not its temporal structure – neither its speed, nor its temporal grouping. This dissociation is counterintuitive, however, we propose that temporal accuracy can be dissociated from timing in CQ models. In our model (Figure 5 and Methods), we assume that positional associations of the items in the sequence (positional context and parallel planning layer) are determined by the respective sequence cue, and the corresponding start-state of the cued sequence becomes gradually activated. Crucially, we show that changing the width of the receptive field for each position (Figure 5a) affects the activation gradient of action items during sequence planning (Figure 5b). Specifically, our model demonstrates that narrowing this positional tuning will cause a steeper relative activation gradient at the end of sequence preparation, with actions in later positions being progressively less activated. Conversely, wider tuning, would broaden the excitation from the positional context to parallel planning layer and lead to smaller relative activation differences between actions at the end of the planning period. Notably, while the positional tuning in CQ models is hypothesized to be acquired through exposure (Hebbian learning) [46], we assume that it is dynamically established throughout the preparation period (cf. Methods section). Thus, the width of the positional tuning of individual actions in the parallel planning layer may underly the accuracy of actions, independently of the overall speed and temporal structure of sequences.

**Figure 5.**
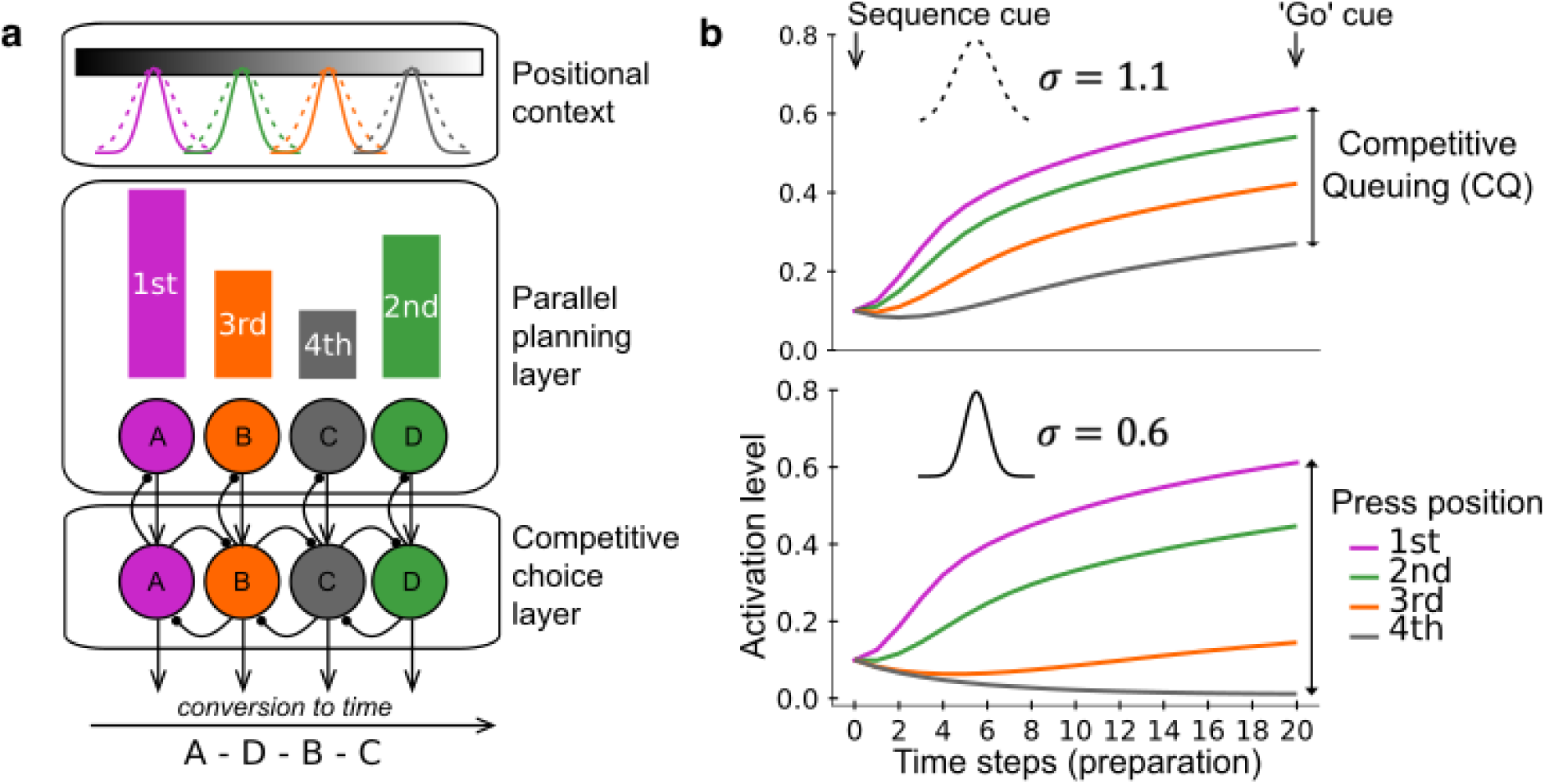
Competitive queuing (CQ) model and the role of positional tuning in sequence preparation. **a**. The Parallel planning and Competitive choice layers of the CQ model contain nodes representing possible sequence items, such as finger presses A, B, C and D. When learning a sequence, connections are formed from sequentially activated nodes in the Positional context layer to item nodes in the Parallel planning layer as each is activated in turn. Crucially, the current model incorporates a positional tuning of the nodes. The receptive field of this positional tuning has a tuning (variance) parameter *σ*, controlling the model’s sensitivity to positional differences. This tuning curve may be acquired though training (variability of instructive stimulus and training exposure), as well as reflect an intrinsic variability of each participant (sensory or motor variability). **b**. The tuning width of the receptive field determines, inversely, the spread of the competitive activations of corresponding action items following the *Sequence* cue, with a narrow or sharper tuning producing more pronounced CQ during preparation. Time steps represent linear arbitrary values.

Although our empirical data and CQ model do not support the integration of the timing signal with action order before sequence execution, they do not exclude the presence of a separate preparation process for the speed and timing of the respective sequence, which may take place concurrently or at different time points during preparation [47–49]. In previous work, we proposed a drift-diffusion based model which contains input from separate modules that activate action order and timing.[50] This model was based on behavioural sequence learning data demonstrating that sequence timing is encoded independently of the action order, but requires multiplicative, rather than additive integration with each action. This enables previously learnt sequence timing to be transferred to new sequences, but only after the action order has been acquired, reconciling previous experimental findings [51–55]. Most recently Zeid and Bullock proposed how such plans might be generated in the context of CQ models [52], proposing that two CQ modules could operate in parallel – one controlling the item order and the other controlling the sequence of inter-onset- intervals that define a rhythmic pattern, including separate parallel planning and competitive choice layers. While this model is in line with neurophysiological and imaging evidence for a separate control of timing for sequence generation [51,56–60], empirical data for a dedicated CQ process for temporal intervals is lacking.

## Conclusions

In sum, our findings indicate that the graded relative activation state during a brief period of retrieval and planning reflects the subsequent action order and correlates with the individual’s sequence fluency and accuracy. It appears to be invariant to the exact timing of the sequence, but is instead bound to the precision of the positional tuning. In contrast to neurophysiological approaches involving advanced neural pattern analysis [11,14], a simple behavioural paradigm could provide a straightforward and cost-effective proxy to assess the state of action preparation across trials in individual participants. This behavioural readout may help advance our understanding of the neural processes associated with disorders affecting the fluent production motor sequences, such as stuttering, dyspraxia, and task-dependent dystonia [61–65].

## Methods

### Participants

Data were collected from a total of 55 right-handed University students (Experiment 1: *N* = 19, 11 females; *M* = 24.2 years, *SD* = 4.1; Experiment 2: *N*=18, 11 females; *M* = 24.2 years, *SD* = 4.5; Experiment 3: *N* = 18, 9 females; *M* = 20.8 years, *SD* = 2.4). Four additional participants were tested, but excluded from analysis based on their sequence production error rate (cf. Participant exclusion criteria). They were hypothesis-naive and had no previous exposure in performing a similar experimental task. All participants had normal or corrected-to-normal vision and reported no history of neurological or psychiatric disorders or hearing problems. Handedness was evaluated through the online Handedness Questionnaire (http://www.brainmapping.org/shared/Edinburgh.php) adapted from the Edinburgh Handedness Inventory [66] (Experiment 1, *M* = 88.4, *SD* = 9.4; Experiment 2, *M* = 90.6, *SD* = 9.7; Experiment 3, *M* = 90, *SD* = 11.8). All participants provided written informed consent before participation and were debriefed after completing the study. They were compensated either monetarily or with course credits at the end of the experiment. All procedures were approved by the Bangor University School of Psychology Research Ethics Committee (Ethics Review Board Approval Code 2017-16100-A14320).

### Participant exclusion criteria

Mean finger and temporal interval error rate during sequence production in the test phase (Day 3) above three standard deviations of the group mean performance was considered as outlier performance, in each experiment separately. This was to ensure that participants reached a comparable skill level in sequence performance from memory and to have sufficient number of trials for RT analysis per participant, which included correct trials only. We set this blindly to the individual *Probe* trial performance to ensure that data exclusion was independent of the data analysed to to test our main hypotheses. This resulted in the exclusion of data from one participant in Experiment 1 who showed 53.1% finger error in the short delay, 54.7% in the intermediate delay and 53.9% in the long preparation duration conditions. Two participants’ data sets were removed from Experiment 2, one with 25% finger error in the slow timing and 18.8% in the irregular timing conditions, and the other with 44.5% in the fast timing conditions. The data of one participant was excluded from Experiment 3 due to 12.5% finger error in the fast timing condition. No outlier performance was found for temporal interval production in any condition of each experiment according to the above criteria. Overall, the data of 19 participants were analyzed for Experiment 1, 18 participants for Experiment 2, and 18 participants for Experiment 3.

### Apparatus

For all three experiments participants were seated in a quiet room in front of a 19-inch LCD monitor (LG Flatron L1953HR, 1280 x 1024 pixels, refresh rate 60Hz), wearing headphones for noise isolation. All instructions about when each block began, visual stimuli and feedback were controlled by Cogent 2000 (v1.29) (http://www.vislab.ucl.ac.uk/cogent.php) through a custom-written MATLAB program (v9.2 R2017a, The MathWorks, Inc., Natick, Massachusetts, United States) and projected on to the LCD screen with inter-stimulus-intervals calculated in refresh rates to ensure precise stimulus timing. In Experiments 1 and 2, a customized foam channel was attached to the outer-half surface of the table to stabilize the cable of a Pyka 5- button fiber optic response device (Current Designs). A thin anti-slip black mat was placed underneath the response device to prevent sliding during the task. The response device was positioned horizontally and adjusted accordingly for each participant to ensure good control over the target buttons as well as arm and wrist comfort. Participants were instructed to place the right index, middle, ring and little fingers on the respective target buttons of the device. Experiment 3 used an identical experimental set-up with the exception that responses were recorded using a computer keyboard and participants were instructed to place their right thumb in addition to the rest of the right-hand fingers on the designated keyboard keys. For hand stabilization and comfort their wrist was positioned on a wrist rest.

### Behavioural task and design

In Experiments 1 and 2, the task involved the recording of sequential and single button presses produced with the four fingers (index, middle, ring and little) of the right hand on a response device while performing a visually cued motor learning task adapted from Kornysheva et al. [11]. Experiment 3 additionally required single presses with the thumb. Participants were trained to associate a visual cue (an abstract fractal shape, henceforth *Sequence* cue) with a specific a four-element finger sequence produced with a specific timing. In all experiments, the paradigm employed two main trial types: sequence and single-press (*Probe*) trials. *Sequence* trials were further divided into visually instructed and memory-guided trials. Instructed trials involved the presentation of four visual digit cues (index, middle, ring and little) at specified intervals comprising a unique target sequence. These were only used during training in the first two days, and during two refresher blocks on the third day. The test phase on the third day only involved sequence production without visual guidance (memory-guided trials, Supplementary Figure 3). *Probe* trials involved the production of only one visual digit cue (*Probe* cue) corresponding to one of the serial positions in the target sequence (Figure 1b).

#### Experiment 1

All participants were trained in producing two four-element target sequences comprising two different finger order types (F1, F2) with isochronous temporal intervals of 800 ms between presses (T1). Two additional finger order types (F3, F4) of the same temporal sequence (T1) served as practice sequences to impose familiarization with the task. Four additional finger order types (F5, F6, F7, F8) with isochronous intervals of 800 ms (T1) were used to evaluate sequence-specific learning in a visually cued task alongside the target sequences, immediately before and after the training phase. The data from this control task is not presented here, as the current work evaluates the preparation and performance during trials involving production from memory. As a result, the experiment employed a total of eight unique *Sequence* cues associated with eight finger sequences. The sequences were randomly generated offline through a custom-written MATLAB code for each participant. Specifically, the sequence generation process produced sequences for each participant randomly excluding sequences with ascending and descending digit triplets. The trained sequences started with different digits.

All trial types started with a *Sequence* cue. The *Sequence* cue had a fixed duration of 400ms followed by a fixation cross, the latency of which varied depending on the delay period from *Sequence* cue onset to *Go* cue. The resultant short (500 ms), intermediate (1000 ms), and long (1500 ms) delay periods following the *Sequence* cue comprised the three preparation duration conditions employed in the task. After the delay period, a black hand stimulus appeared as the *Go* cue. In an instructed trial, the *Go* cue was presented on a grey background for 2400 ms, guiding the participants throughout the execution of the sequence by sequentially displaying a small white circle on the digits of the hand stimulus. This acted as a visual digit cue appearing sequentially on each of the four digits, with the time intervals between the digit cues forming the target temporal structure of the sequence (T1) and defining its duration of 2400 ms. To achieve finger and temporal accuracy during training, participants were asked to press the correct target buttons in synchrony with the digit cues until the completion of the sequence, with the aim to progress towards synchronization with the target timing. As the first digit cue of a sequence appeared at the same time as the *Go* cue, immediate initiation of the sequence was emphasized in the instructions.

In memory-guided trials, a green rectangle was used as a background for the *Go* cue, remaining on the screen for 2400 ms. Memory-guided trials featured the *Go* cue without the appearance of digit cues, requiring participants to produce the upcoming target sequence from memory. In these trials, participants were instructed to initiate the sequence as quickly as possible and produce the sequence according to its target finger and temporal structure (i.e. F1T1, F2T1).

In probe trials, the *Go* cue was displayed for 1000 ms on a grey background with a digit cue presented on one digit (*Probe* cue), prompting a single press of the corresponding target button. Here, the instructions were to respond to the *Probe* cue as fast and accurately as possible. Participants were encouraged to avoid premature responses (before the *Go* cue) in all trial types.

Following the *Go* cue, a fixation cross (1000 ms) and, subsequently, feedback (1000 ms) were presented on the screen. The duration of a sequence trial including feedback was 5.4 s, while a probe trial had a duration of 4 s. The inter-trial-interval (ITI) was fixed at 800 ms. The experiment consisted of two 90min long training (Days 1 and 2) and a test (Day 3) sessions taking place over three consecutive days. Day 1 commenced with a practice block which involved two instructed and two memory-guided *Sequence* trials for each of the target finger sequences as well as two random probe trials, all randomly combined with the three delays. Over the three days, participants serially underwent a pre-training (2 blocks), a training (36 blocks), a post-training (2 blocks) and a test phase (2 refresher training blocks + 16 test blocks) completing a total of 58 blocks. Participants were naïve as to the structure of the gradual transition from the training through to the testphase and which block type they were administered. The pre- and post-training blocks consisted of 24 instructed trials each; each block was 2.48 min long and contained randomized mixed repetitions of the two target and four control sequences matched equally with the delay conditions. The training phase was organized in three stages: 12 blocks of 288 instructed and 72 probe trials (stage A, 80% instructed and 20% probe trials in each block), 12 blocks of 144 instructed, 144 from memory and 72 probe trials (stage B, 40% for each sequence type and 20% probe trials in each block), and 12 blocks of 288 memory-guided and 72 probe trials (stage C, 80% memory and 20% probe trials in each block). A training block (3 min long) of either stage consisted of 30 trials. On each block there was a stable 20% occurrence of probe trials (6 in each block) comprising a total of 216 probes throughout the training blocks. Distribution of probe trials in this phase was determined by the minimum number of trials possible, namely 24 (2 sequences × 3 delays × 4 probe digits), and the block repeats. Eventually each probe digit occurred 18 times in each training stage. All 40 blocks were evenly assigned to the study sessions such that from Day 1 through the end of Day 2 participants had completed the training and the post-training synchronization task. The testphase (Day 3) started with two refresher training blocks of mixed type and immediately progressed to 16 blocks of 48 trials each, in which 24 memory-guided and 24 probe trials were randomly presented. Duration of a test block was 4.4 min. The two trained sequences used in the memory- guided trials were matched to the three delay conditions with each combination being repeated four times within the block. This gave a total of 128 memory-guided trials per delay condition, across blocks. In probe trials, each probe digit was combined with the three delay conditions resulting in 32 trials per digit per delay condition. The testphase had a total of 768 trials (384 memory-guided sequences and 384 probes). Overall, the participants underwent 2004 trials excluding the practice trials.

#### Experiment 2

Procedures for Experiment 2 were identical to Experiment 1 except that the preparation period was fixed at 1500 ms and participants were trained in associating three unique *Sequence* cues with one finger sequence (F1) to be performed with three target temporal structures (T1, T2, T3) or IPIs: slow (T1, 800- 800-800 ms), fast (T2, 400-400-400 ms) and irregular (T3, 400-1600-400 ms), forming respective target sequence durations of 2400 ms, 1200 ms, and 2400ms. The trial followed the same structure as in Experiment 1, but the *Go* cue remained on the screen for 3000 ms in a sequence trial and for 1000 in a probe trial. This was followed by a fixation cross (1000 ms) and feedback (1000 ms) with a varying ITI of 500, 900 and 1300 ms. As a result, a sequence trial was 6.5 min long and a probe trial 4.5 min long. The participants underwent the same structure of training and testsessions as in Experiment 1. Similarly, we conducted a synchronization task, in a pre-post design. Here, three additional control timing conditions were used to test for temporal transfer (trained timing conditions) with the target timing patterns combined with a different finger sequence. Overall, in this experiment participants were exposed to 15 unique temporally structured sequences associated with their respective *Sequence* cue and completed 2016 trials over 58 blocks.

#### Experiment 3

The training/testprocedures, trial structure, and the pre/post-training synchronization task in Experiment 3 were identical to those of Experiment 2, except that probe trials would additionally cue the thumb. This served as a control condition to obtain reaction times and error rates for unplanned responses as thumb presses were not part of any learnt finger sequence. Across each training stage, there were 60 probe trials, while the testphase (30 blocks × 26 trials) contained 360 memory-guided *Sequence* trials (120 trials per timing condition), 360 probe trials (30 trials per digit per timing condition), and 60 thumb probe trials (20 trials per timing condition). Overall, participants completed 1990 trials over 72 blocks, excluding the practice block.

#### Feedback

In all experiments, a points system was designed to reward fast initiation and accurate performance, and avoid any drift in the motor production from memory. After each sequence trial, feedback was presented on the screen for 1000 ms in the form of points (0-10) based on three performance criteria: reaction time (RT) to assess sequence initiation, percentage of deviation from the target temporal intervals of the sequence, and finger press accuracy. Points gained from the RT component of the sequence, i.e. response from *Go* cue to the first press, were defined by tolerance RT windows of 0-200, 200-360, 360-480, 480-560, 560-600 ms resulting in 5, 4, 3, 2 and 1 points, respectively. For late (> 600) responses, 0 points were given. A schematic feedback provided information on both finger accuracy and temporal sequence accuracy performance. An ‘*x*’ or a ‘*-*’ symbol was shown for every correct or incorrect press, respectively. Temporal errors were calculated after each trial as deviations of press from target timing in percent of the target interval to account for the scalar variability of timing[67,68]. Thresholds for mean absolute percentage deviation across all correct presses were set at 10, 20, 30, 40 and 50 percent assigning 5, 4, 3, 2 and 1 points, respectively. Timing interval deviation > 50% resulted in 0 points. If a press was performed too early the respective symbol was displayed below the midline, while for a late response it was displayed above. This applied only to the second, third and fourth presses of the sequence, whilst the first symbol reflecting the first press was always positioned on the midline, representing the starting point of the sequence. Deviation from target onset (presented or assumed) rather than interval timing encouraged participants to synchronize with the visually cued sequences during training, however, may have contributed to a tendency to compress the overall sequence length during trials produced from memory.

Participants were instructed to adjust their performance by keeping the crosses as close to the midline as possible. If at least one incorrect press or an incorrect number of presses was recorded (< 4 or > 4), 0 points were given on that trial. The points on each trial were displayed above the schematic feedback, and were the sum of the RT, interval deviation and finger accuracy points. The feedback following a probe trial displayed only points (0-5) gained based on RT and finger press accuracy utilizing the same tolerance windows as described above for assessing sequence initiation RT. In the case of an incorrect press or incorrect number of presses (< 1 or > 1), 0 points were given regardless of the RT length. To incentivize the participants to gain as many points as possible on each trial, we offered an extra monetary reward (10£) to those two with the highest total points.

### Data analysis

Data analyses were performed using custom written code in Matlab (v9.2 R2017a, The MathWorks, Inc., Natick, Massachusetts, United States) and SPSS version 22.0 (IBM Corp., Armonk, N.Y., USA). Median reaction time (RT; correct trials only) and mean error rates for each *Probe* position and condition were calculated relative to the 1st position in each participant and condition (RT and error rate increase in %).

Repeated measures ANOVAs were undertaken for RT and error rate in *Probe* trials, and for inter-press-intervals (IPI), temporal error, finger error, and sequence initiation RT in *Sequence* trials produced from memory. Planned contrast analyses for the main and interaction terms of interest in each ANOVA model involved user-defined orthogonal contrasts. To evaluate the RT and error rate increase for the control action (Experiment 3), we used two-tailed paired-samples t-tests (control *vs* 4th position).

Mean relative increase between adjacent positions (1st to 2nd, 2nd to 3rd, 3rd to 4th) for RT (CQ RT increase) and press error rate (CQ error increase) in *Probe* trials were taken as a measure for the strength of the competitive action gradient during preparation. Using the group data (N = 55), we conducted six planned one-tailed Pearson correlations between the CQ strength derived from RT and error increase, respectively, and each of the sequence production measures.

### CQ model of sequence preparation

To our knowledge, no CQ model has previously been applied to response preparation. While models differ somewhat with respect to how sequence position is represented, they all require some form of “start state”, which has stronger links to items or responses that should occur earlier in the sequence [5,8,69,70]. In the implementation, we use the Start-End CQ model [16], but we expect that other CQ models with distinct start states would behave similarly if the same assumptions regarding preparation were added to them.

The model makes the following assumptions: Each learned sequence is hierarchically organized, with a “sequence node (or nodes)” linking to, and activating, the responses which make up the sequence. The sequence node activates the position codes (associations) of the items in the sequence. Following the *Sequence* cue, the start-state of the cued sequence becomes gradually activated. The start-state of the intended sequence produces an activation gradient over the planned responses based on its strength of association with them. The additions to the published CQ model of Houghton [16] consist of: (i) the gradual increase of the start state activation, (Equation 1); (ii) the damping of response activations prior to the *Go* cue (Equation 3); and inhibition of the competitive response selection process, also prior to the *Go* cue. The model’s activation of planned responses during preparation is shown in Figure 5.

#### Sequence node activation

Following presentation of the *Sequence* cue, the associated sequence node *s*_*j*_ becomes gradually activated. We implement a simple linear “ramp”,

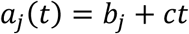

where *t* is (discrete) time (since cue presentation), *b*_*j*_ is the baseline activation, and *c* is the rate of increase. The activation of the start-state *S*_*j*_ retrieved by the learned sequence *s*_*j*_ follows this activation,

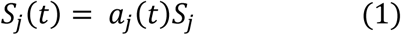

*S*_*j*_ (without a time index) is the asymptotic (i.e., stored) value for the sequence, set here to unity. The effect of this is simply that the start-state gradually increases its activation following the cue.

#### Input to response nodes

Activation spreads from the start-state *S*_*j*_(*t*) to finger responses via its positional associations *W*_*j*_ (a matrix) with the response tokens[16]. The input from sequence *s*_*j*_ to its associated actions is given by,

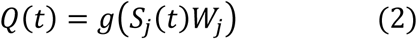

Here *Q*(*t*) (a matrix) represents the state of the queue of response tokens. *W*_*j*_ encodes the positional weights from the sequence level to the responses in *s*_*j*_, and the product *S*_*j*_(*t*)*W*_*j*_ computes the differences between the start state of the sequence (*S*_*j*_(*t*)) and the position codes *W*_*j*_ of the sequence items (represented using the phase code of Houghton [16]).

Finally, the function *g* represents a Gaussian receptive field, or positional tuning curve, applied element-wise to the signals coming from the matrix *W*_*j*_. The receptive field is sensitive to the difference between the state of the start-signal *S*_*j*_ and the position codes in *W*_*j*_, peaking when they are identical (see Houghton, 2018, Equation 1). The receptive field has a tuning (variance) parameter *σ*, controlling the model’s sensitivity to positional differences (Figure 5a).

#### Finger response activation

Each finger response in the sequence *s*_*j*_ becomes gradually active during preparation, due to the increasing activation of the sequence’s start state (Equation 1), sending an increasing degree of activation to the responses in the sequence, modulated by the similarity of the response’s position code to the start state (Equation 2). For a finger response *F*_*k,j*_ (i.e., *k*th finger in sequence *j*), its activation during preparation is, in discrete time form,

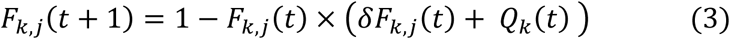

In the second term on the right, *δ* controls the decay rate (of the current activation level), and *Q*_*k*_(*t*) is the input to finger *F*_*k*_ given by Equation 2. Note that if the latter equals 0, then response activation spontaneously decays due to the decay term, *δ* < 1.

The first term on the right, 1 − *F*_*k,j*_(*t*), acts as “damping” factor; it becomes smaller as the response activation increases towards a ceiling of 1. This prevents activations growing without bound as the preparation interval increases (Figure 5b). It is proposed that on detection of the *Go* signal, this damping term ceases to act, permitting activations to rapidly increase, and initiating the competitive response selection process intrinsic to CQ models.

## Acknowledgements

The authors wish to thank Tom Hartley, Ken Valyear and Simon Watt for helpful comments on the study.

