## Supplementary Figures for "Competitive state of actions during planning predicts sequence execution accuracy"

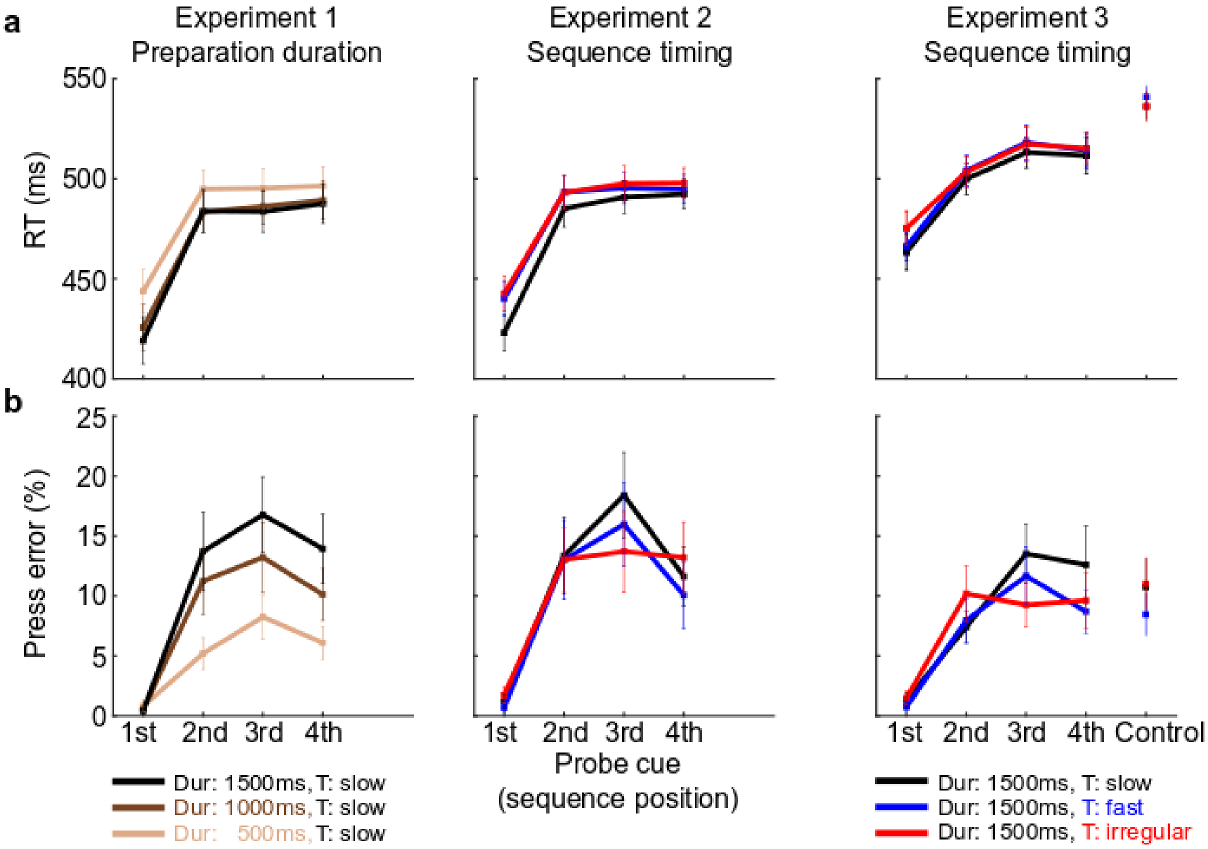


**Supplementary Figure 1 | Raw reaction time (RT) and press error rates of single actions (*Probe* trials).** Complementary data to RT and Error increase in Figure 3. **a.** Mean RT increased with sequence position number up to the 4th position in Experiment 1 (2nd position *vs* 1st position, $F$ (1, 18) = 83.165, $p$ < .001, $\eta p^2$ = .822; 3rd position *vs* 2nd position, $F$ (1, 18) = 66.079, $p$ < .001, $\eta p^2$ = .786; 4th position *vs* 3rd position, $F$ (1, 18) = 26.187, $p$ < .001, $\eta p^2$ = .593) and in Experiment 2 (2nd position *vs* 1st position, $F$ (1, 17) = 60.271, $p$ < .001, $\eta p^2$ = .780; 3rd position *vs* 2nd position, $F$ (1, 17) = 34.053, $p$ = .001, $\eta p^2$ = .667; 4th position *vs* 3rd position, $F$ (1, 17) = 28.949, $p$ < .001, $\eta p^2$ = .630) and up to the 3rd position in Experiment 3 with the 4th position performing faster (2nd position *vs* 1st position, $F$ (1, 17) = 66.416, $p$ < .001, $\eta p^2$ = .796; 3rd position *vs* 2nd position, $F$ (1, 17) = 69.965, $p$ < .001, $\eta p^2$ = .805; 4th position *vs* 3rd position, $F$ (1, 17) = 13.417, $p$ = .002, $\eta p^2$ = .441). Preparation duration did not affect RT as in the normalized RT slowing data (Experiment 1, $F$ (6, 108) = 2.069, $p$ = .063, $\eta p^2$ = .103). The absent significant Position x Delay interaction was driven by a non-significant difference between the 3rd *vs* 2nd positions contrasts at the long *vs* short preparation duration (Experiment 1, $F$ (1, 17) = 3.542, $p$ = .076, $\eta p^2$ = .164). Similar to the normalized RT slowing findings, raw RT did not change by position when preparing sequences of different timing structure (Experiment 2, $F$ (3.267, 55.541) = 2.302, $p$ = .082, $\eta p^2$ = .119, Greenhouse-Geisser corrected, $\chi 2$ (20) = 42.608, $p$ = .003; Experiment 3, $F$ (3.870, 65.786) = .975, $p$ = .426, $\eta p^2$ = .054, Greenhouse-Geisser corrected, $\chi 2$ (20) = 34.057, $p$ = .028). The mean RT of the control action was significantly slower than the 4th position indicating less activation in the preparation phase (Experiment 3, $t$ (17) = 3.044, $p$ = .007, $d$ = .856, two-tailed). **b.** Press error rate equally showed a graded increase up to the 3rd position (Experiment 1, 2nd position *vs* 1st position, $F$ (1, 17) = 29.613, $p$ < .001, $\eta p^2$ = .622; 3rd position *vs* 2nd position, $F$ (1, 17) = 25.863, $p$ < .001, $\eta p^2$ = .590; 4th position *vs* 3rd position, $F$ (1, 17) = 5.074, $p$ = .037, $\eta p^2$ = .220; Experiment 2, 2nd position *vs* 1st position, $F$ (1, 17) = 23.940, $p$ < .001, $\eta p^2$ = .585; 3rd position *vs* 2nd position, $F$ (1, 17) = 15.676, $p$ = .001, $\eta p^2$ = .480; 4th position *vs* 3rd position, $F$ (1, 17) = 1.314, $p$ = .268, $\eta p^2$ = .072). This increase was enhanced by longer preparation duration (Position x Delay, Experiment 1, $F$ (4.135, 74.433) = 3.860, $p$ = .006, $\eta p^2$ = .177, Greenhouse-Geisser corrected, $\chi 2$ (20) = 33.864, $p$ = .030; Special contrasts: 2nd position *vs* 1st position at long *vs* short delay, $F$ (1, 18) = 11.015, $p$ = .004, $\eta p^2$ = .380, 2nd position *vs* 1st position at long *vs* intermediate delay, $F$ (1, 18) = 2.968, $p$ = .102, $\eta p^2$ = .142, 3rd position *vs* 2nd position at long *vs* short delay, $F$ (1, 18) = .000, $p$ = .996, $\eta p^2$ = .000, 3rd position *vs* 2nd position at long *vs* intermediate delay, $F$ (1, 18) = .290, $p$ = .597, $\eta p^2$ = .016, 4th position *vs* 3rd position at long *vs* short delay, $F$ (1, 18) = .120, $p$ = .733, $\eta p^2$ = .007, 4th position *vs* 3rd position at long *vs* intermediate delay, $F$ (1, 18) = .011, $p$ = .918, $\eta p^2$ = .001) but not affected by sequence timing (Position x Timing, Experiment 2, $F$ (6, 102) = 1.491, $p$ = .189, $\eta p^2$ = .081; Experiment 3, $F$ (3.823, 64.987) = 1.876, $p$ = .128, $\eta p^2$ = .099, Greenhouse-Geisser corrected, $\chi 2$ (20) = 41.603, $p$ = .004). Error rate of control action was not significantly greater than the 4th position (Experiment 3, $t$ (17) = -.938, $p$ = .361, $d$ = .316, two-tailed).

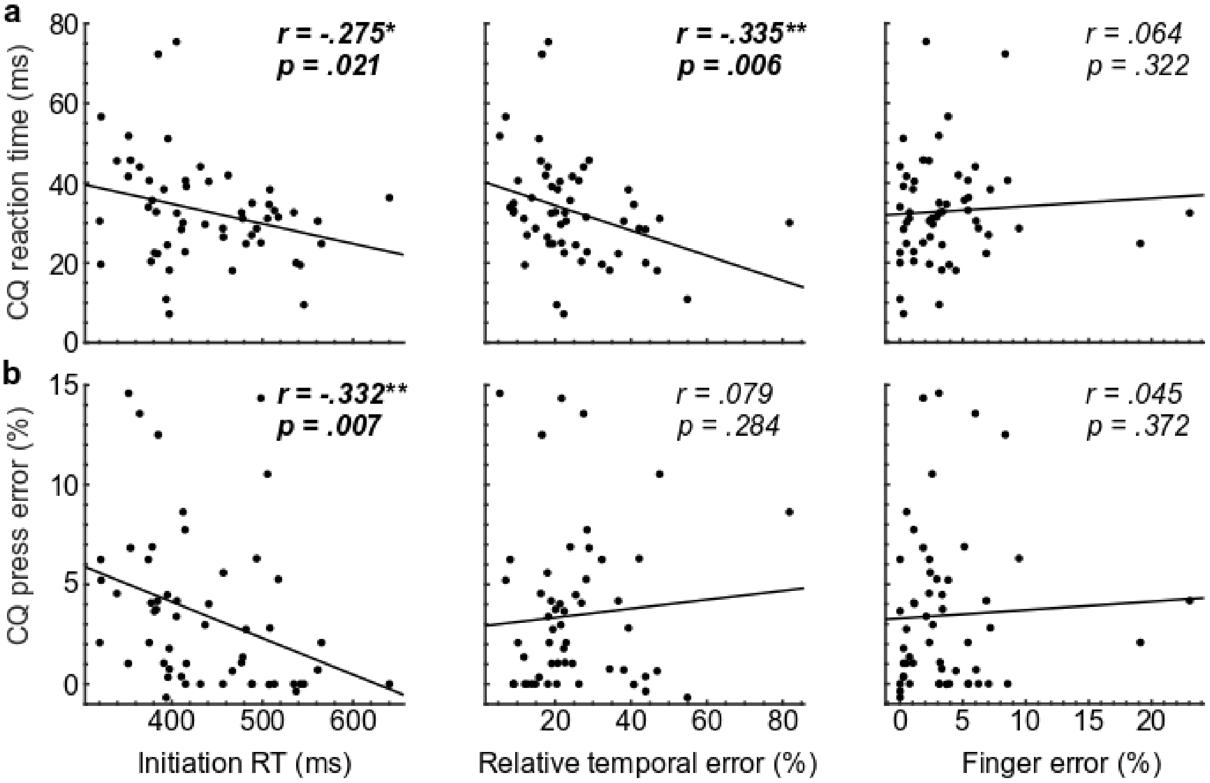


**Supplementary Figure 2 | Raw CQ gradient during preparation correlates with sequence production.** Mean reaction time (RT) in ms and percent press error during preparation (probe trials) were used to calculate the distance between adjacent positions (1st minus 2nd and so on), reflecting the CQ of constituent actions before production. Results replicate the correlation coefficients found for the CQ RT and error increase measures. **a.** A stronger CQ RT gradient, indicating a larger (steeper) CQ during preparation, correlated with faster sequence initiation and less relative temporal errors during production. **b.** The CQ error gradient similarly correlated with faster sequence initiation but not relative temporal accuracy. Neither measure was associated with decreased finger error rate (**a, b**). All correlations are one-tailed.

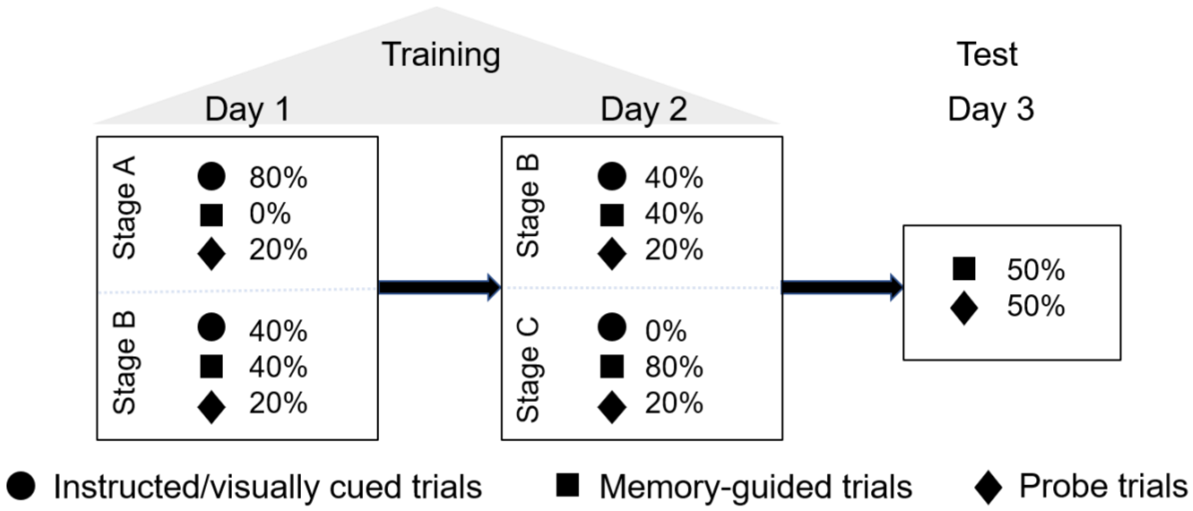


**Supplementary Figure 3 | Two-day training and testing procedure.** The first two days integrated the three training stages. Participants gradually progressed from entirely instructed sequence production trials (stage A) to blocks of mixed trials (stage B) and, finally, to producing the target sequences from memory during the last stage of the training (stage C). All training stages incorporated a fixed percentage of probe trials, randomized in each block, to ensure a degree of familiarity with single-press Probe cues. In the testing phase (Day 3), participants underwent an equal number of memory-guided sequence production trials and probe trials.
